# “Visual” Cortices of Congenitally Blind Adults Respond to Executive Demands Authors

**DOI:** 10.1101/390450

**Authors:** Rita E. Loiotile, Marina Bedny

## Abstract

How functionally flexible is human cortex? In congenitally blind individuals, “visual” cortices are active during auditory and tactile tasks. The cognitive role of these responses and the underlying mechanisms remain uncertain. A dominant view is that, in blindness, “visual” cortices process information from low-level auditory and somatosensory systems. An alternative hypothesis is that higher-cognitive fronto-parietal systems take over “visual” cortices. We report that, in congenitally blind individuals, right-lateralized “visual” cortex responds to executive-load in a go/no-go task. These right-lateralized occipital cortices of blind, but not sighted, individuals mirrored the executive-function pattern observed in fronto-parietal systems. In blindness, the same “visual” cortex area, at rest, also increases its synchronization with prefrontal executive control regions and decreases its synchronization with auditory and sensorimotor cortices. These results support the hypothesis of top-down fronto-parietal takeover of “visual” cortices, and suggest that human cortex is highly flexible at birth.

## Introduction

Across individuals, different cognitive functions are implemented in consistent cortical locations, each of which has a distinctive cyto-architecture and inter-regional connectivity profile. This systematic relationship between structure and function suggests that intrinsic physiology tightly constrains each cortical region to implement particular cognitive operations. Contrary this idea, studies of sensory loss, such as in blindness and deafness, demonstrate that experience can modify the structure to function mapping. In blind individuals, retinotopic “visual” cortices respond to auditory and tactile stimuli (Sadato et al., 1996; Wanet-Defalque et al., 1988), and in deaf individuals, auditory cortices respond to visual and tactile stimuli (Finney, Fine, & Dobkins, 2001; Levänen, Jousmäki, & Hari, 1998). Studies of sensory loss demonstrate that experience can change the sensory modality to which a cortical area responds. However, the extent to which cortical regions truly change their function, even in cross-modal plasticity remains debated (Amedi, Hofstetter, Maidenbaum, & Heimler, 2017; Bavelier & Neville, 2002; Bedny, 2017).

One view is that sensory cortices preserve their original cognitive operation, even in cases of cross-modal plasticity (Amedi et al., 2017; Cecchetti, Kupers, Ptito, Pietrini, & Ricciardi, 2016; Meredith et al., 2011; Pascual-Leone & Hamilton, 2001; Renier et al., 2010; Striem-Amit, Dakwar, Reich, & Amedi, 2011). According to the metamodal hypothesis, in blindness, “visual” cortices continue to perform vision-like functions, but over input from audition and touch. Consistent with this idea, dorsal occipital areas that are part of the visual “where” pathway in sighted individuals become active during sound localization in blind individuals (Collignon, Vandewalle, & Voss, 2011; Gougoux, Zatorre, Lassonde, Voss, & Lepore, 2005; Wanet-Defalque et al., 1988). Analogously, it has been suggested that retinotopic areas typically involved in fine-grained visuospatial discrimination become involved in finegrained somatosensory discriminations, such as texture perception, in blindness (Merabet et al., 2004; Sadato et al., 1996; Sathian & Stilla, 2010). There is also evidence that in deaf cats, auditory areas involved in peripheral sound localization are recruited during localization of peripheral visual stimuli (Meredith et al., 2011). In these instances of cross-modal plasticity, sensory cortices appear to preserve their underlying cognitive operation, even when the sensory modality, to which they respond, changes. One interpretation of these findings is that while the preferred sensory modality is malleable, the cognitive operation itself (e.g. spatial localization) is specified by intrinsic physiology (Pascual-Leone & Hamilton, 2001).

An alternative possibility is that cortices are capable of drastically altering their function based on early experience (Bedny, 2017). Evidence for this idea comes from studies of blindness which demonstrate that “visual” cortices become responsive to language. In blindness, retinotopic “visual” areas, including V1, become sensitive to meaning and grammar. In blind, but not sighted, individuals, occipital cortices respond more to words than meaningless sounds, more to sentences than unconnected lists of words, and more to grammatically complex than grammatically simple sentences (Bedny, Pascual-Leone, Dodell-Feder, Fedorenko, & Saxe, 2011; Lane, Kanjlia, Omaki, & Bedny, 2015; Röder, Stock, Bien, Neville, & Rösler, 2002). Furthermore, language-responsive “visual” cortices become correlated at rest with prefrontal language regions (Bedny et al., 2011). Since language and vision are cognitively and evolutionarily distinct, these observations challenge the idea that cortical areas have fixed functions, even metamodal ones.

There are, however, ways to reconcile findings of language repurposing in “visual” cortices of blind individuals with the idea that cortical areas have fixed functions. One possibility is that the occipital cortices are specifically predisposed for both vision and language. For example, visual scene perception and sentence processing could share an underlying cognitive operation, such as hierarchical structure building. Another possibility is that Braille bootstraps the “visual” cortices into language processing (Bavelier & Neville, 2002). Braille recognition is similar to vision in that both involve fine-grained spatial discrimination; language processing could invade the visual cortices as a secondary consequence of Braille learning. Therefore, findings from language could be reconciled with the idea that visual areas preserve their underlying functions in blindness.

A key open question, therefore, is whether language is the only higher-cognitive function assumed by “visual” cortices in blindness. If so, language encroachment into the “visual” system may be a special case of metamodality. An alternative possibility is that language encroachment into the visual system is part of a broader phenomenon, whereby the functional specialization of deafferented “visual” cortices is driven by top-down anatomical inputs from prefrontal, parietal, and temporal networks (Bedny, 2017). In sighted individuals, these projections enable communication between visual and higher cognitive systems, as well as between vision and other sensory modalities (Bressler, Tang, Sylvester, Shulman, & Corbetta, 2008; Corbetta & Shulman, 2002; Egner & Hirsch, 2005; Gilbert & Li, 2013; Kastner & Ungerleider, 2000; Tong, 2003). According to the higher-cognitive takeover hypothesis, in blindness, the absence of bottom-up input from the lateral geniculate nucleus (LGN) enables top-down projections to colonize the visual system for an array of verbal and non-verbal higher-cognitive operations (Bedny, 2017).

Preliminary evidence for the higher-cognitive takeover hypothesis comes from a recent study of mathematical processing in blindness. Dorsal retinotopic “visual” areas are active when congenitally blind individuals solve spoken math equations (e.g. 17-4=X), more so than when blind participants listen to non-mathematical sentences, and the amount of activity scales with equation difficulty (Kanjlia, Lane, Feigenson, & Bedny, 2016). These math-responsive “visual” regions are differentially localized within occipital cortices from sentence-responsive regions and show a distinctive functional connectivity profile with the fronto-parietal number network (Kanjlia et al., 2016). Furthermore, even at rest, their activity is correlated with fronto-parietal regions in blind individuals (Kanjlia et al., 2016).

These results provide tentative support for the idea that language is not the only higher-cognitive function found in deafferented visual cortices. However, spoken math equations arguably share important properties with language: they include spoken words, they are symbolic, they involve hierarchical structure, and they can also be written in Braille. An outstanding question is whether “visual” cortices of blind individuals are also involved in entirely non-verbal higher-cognitive functions.

Fronto-parietal executive functions offer a natural test case for answering this question. Executive functions regulate behavior towards task-relevant goals through processes such as selective attention and response selection (Banich, 2009; Diamond, 2013; Miyake, 2000). In sighted individuals, fronto-parietal executive systems modulate activity in visual cortices during visual perception tasks (Desimone & Duncan, 1995; Miller & Cohen, 2001; Moran & Desimone, 1985). This is accomplished via known anatomical projections (in primates) to the visual system from polymodal parietal and, to a lesser degree, frontal cortices (Anderson, Kennedy, & Martin, 2011; Martino, Brogna, Robles, Vergani, & Duffau, 2010; Maunsell & Van Essen, 1983; Rockland & Ojima, 2003; Selemon & Goldman-Rakic, 1988; Ungerleider, Courtney, & Haxby, 1998; Ungerleider, Galkin, Desimone, & Gattass, 2008; Yeterian, Pandya, Tomaiuolo, & Petrides, 2012). Executive systems are, therefore, likely to constitute a robust input to deafferented “visual” cortices in blindness. The higher-cognitive takeover hypothesis predicts that “visual” cortices of blind individuals take on domain-general executive operations, apart from language processes.

A handful of previous studies are broadly consistent with the idea that “visual” cortices take on non-verbal executive functions in blindness. For example, Park et al. (Park et al., 2011) reported greater “visual” cortex activity during a 2-back than a 0-back control task with tones. Electrophysiological and fMRI studies find that “visual” cortices of blind individuals respond to deviant presentations of tones and tactile stimuli. These responses are thought to reflect attentional, rather than automatic sensory, processes because they occur later and only for attended stimuli (Kujala et al., 1997; Kujala, Alho, et al., 1995a; Kujala, Huotilainen, et al., 1995b; Kujala et al., 2005; Liotti, Ryder, & Woldorff, 1998; Weaver & Stevens, 2007). Another study observed elevated responses in “visual” cortices of blind individuals during the response portion of working memory task, when participants were making a button press (Bedny, Pascual-Leone, Dravida, & Saxe, 2012). These studies provide some evidence that “visual” cortices are sensitive to the higher-cognitive demands of non-verbal tasks. However, the precise cognitive processes performed by “visual” cortices during these tasks remain uncertain and alternative explanations in terms of sensory stimulation have not been ruled out (e.g. (Burton, Sinclair, & Dixit, 2010; Burton, Sinclair, & McLaren, 2004)).

Therefore, the goal of the current study was to test the prediction that regions within the “visual” cortices of blind individuals are incorporated into non-verbal executive function networks. Specifically, we predicted that in blindness a subset of visual cortex would be sensitive to response selection demands in a non-verbal go/no-go task when other factors, such as somatosensory stimulation, are controlled. To test these predictions, congenitally blind and sighted-blindfolded participants performed an auditory go/no-go task with complex non-verbal sounds.

During the go-no/go task, participants made button-presses to some sounds (go trials) and withheld responses to other sounds (no-go trials). Go trials were much more frequent than no-go trials (25% vs. 75%) and participants had to respond quickly (within 900 MS). As a result, the button press becomes pre-potent and must inhibited on no-go trials (Aron, Robbins, & Poldrack, 2014; Garavan, Ross, & Stein, 1999). The increased executive demands of no-go relative to go trials are evidenced both behaviorally and neurally. Participants make more errors of commission (going on no-go trials) than errors of omission (not going on go-trials). Neurally, no-go trials produce elevated activity in right-lateralized fronto-parietal executive function networks among sighted individuals (Barber, Caffo, Pekar, & Mostofsky, 2013; Chikazoe et al., 2008; Garavan et al., 1999; Garavan, Ross, Murphy, Roche, & Stein, 2002; Konishi, Nakajima, Uchida, Sekihara, & Miyashita, 1998; Liddle, Kiehl, & Smith, 2001; Menon, Adleman, White, Glover, & Reiss, 2001; Mostofsky et al., 2003). We predicted that “visual” cortices of blind individuals would respond more to no-go than to go trials, indicating recruitment for non-verbal executive functions and in particular of response selection demands.

Importantly, the current design enables us to distinguish “visual” cortex responses to executive demands from other potentially confounded processes. First, since the current task does not involve language stimuli, “visual” cortex responses are unlikely to be related to language processing. We further predicted that unlike previously observed responses to language in the “visual” cortices, responses to domain-general executive demands would be right-lateralized, similar to responses to executive demands in the fronto-parietal cortices (Aron, 2006; Aron, Robbins, & Poldrack, 2004; Wager et al., 2005). Second, the current task was not spatial; therefore, observed effects are unlikely to reflect vision-like processing. Finally, the design pitted executive demands against low-level sensorimotor demands. If “visual” cortices of blind individuals respond to executive demands, they should be more active during no-go trials. By contrast, if the “visual” cortices respond to sensorimotor demands, they should be more active during go trials, since only the go trials contain a button press and associated tactile feedback. Indeed, previous studies have shown that unlike executive function networks, sensorimotor cortices respond more to go trials than no-go trials (Garavan et al., 1999; Liddle et al., 2001; Mostofsky et al., 2003). Thus, in the current experiment we predicted a double dissociation between activity in sensorimotor cortices and activity in the “visual” cortices of blind individuals.

In the current version of the go/no-go task we also included an intermediate executive demand condition, the infrequent-go. The infrequent-go condition was associated with a distinct sound; it occurred only 25% of the time (like the no-go condition) and required a button press response (unlike the no-go condition). All together there were thus three types of trials: frequent-go (50%), infrequent-go (25%), and no-go (25%). A previous study using a similar design observed an intermediate level of activity for the infrequent-go condition (less activity than no-go but more activity than frequent-go) in prefrontal executive function areas of sighted individuals (Chikazoe et al., 2008). We, therefore, predicted that “visual” cortices of blind individuals would respond most to no-go trials, followed by infrequent-go trials, and least to frequent-go trials.

A second prediction of the current study was that executive-load responsive “visual” areas would become functionally connected at rest with fronto-parietal executive function systems in blindness. To test this prediction, we collected resting state data from a large sample of congenitally blind (n=25) and sighted (n=25) participants. We then asked whether the connectivity of executive-function responsive “visual” cortex is stronger with fronto-parietal executive function networks than with either non-visual sensory-motor areas (early auditory and somatosensory cortices) or language responsive prefrontal cortices. Such a result would support the hypothesis that these “visual” cortex regions are incorporated into the executive system.

In sum, we make four predictions: (1) that the occipital cortices of the blind, but not sighted, group will respond to executive function demands, i.e. most to no-go trials and least to frequent-go trials; (2) that the sensorimotor cortices will display the opposite ordering of responses to the conditions, i.e. most activity for go and least activity for no-go trials, thereby diverging from the executive function profile observed in the blind group’s “visual” cortices; (3) that “visual” cortex responses to executive function demands will be right-lateralized and, thereby, both neuroanatomically dissociable from “visual” cortex responses to language and co-lateralized with fronto-parietal responses to executive function; and (4) that, at rest, executive-function responsive “visual” cortices of blind individuals will show increased functional connectivity to fronto-parietal executive function regions, specifically.

## Results

### Behavioral performance

In both go conditions, participants responded quickly (MS: sighted frequent-go mean=366.77, s.d.=52.80; sighted infrequent-go mean=407.87, s.d.=51.94; blind frequent-go mean=345.70, s.d.=67.92; blind infrequent-go mean=378.26, s.d.=60.74) and made few errors of omission (% correct: sighted frequent-go mean=96.88, s.d.=4.35; sighted infrequent-go mean=95.47, s.d.=4.49; blind frequent-go mean=95.61, s.d.=8.14; blind infrequent-go mean=95.30, s.d.=9.22). Both blind and sighted participants made some errors of commission on no-go trials (% correct: sighted mean=83.28, s.d.=11.17; blind mean=86.88, s.d.=11.82).

Participants in both groups made more errors on no-go than frequent-go or infrequent-go trials (frequent-go vs. no-go sighted t(18)=5.54, p<0.001, blind t(18)=3.40, p=0.003; infrequent-go vs. no-go: sighted t(18)=5.00, p<0.001, blind t(18)=3.16, p=0.005), with no difference between groups (group X condition ANOVA: main effect of condition: F(2,72)=35.08, p<0.001; main effect of group: F(1,36)=0.10, p>0.05, group X condition interaction: F(2,72)=1.48, p=0.235). Frequent and infrequent-go accuracy were different in the sighted, but not the blind, group (sighted t(18)=2.10, p=0.05, blind t(18)=0.63, p>0.5). Differences between the two go conditions were evidenced in response times for both groups: blind and sighted groups were slower to respond on infrequent-go than frequent-go trials (sighted t(18)=7.67, p<0.001, blind t(18)=6.19, p<0.001; group x condition ANOVA: main effect of condition F(1,36)=96.27, p<0.001, main effect of group F(1,36)=1.84, p=0.18, group-by-condition interaction F(1,36)=1.30, p=0.26).

## fMRI Results

### Right-lateralized “visual” cortices of blind individuals responds to executive function, similar to right-lateralized prefrontal cortices (individual subject functional ROI analysis)

In both the sighted and the blind groups, areas of the right prefrontal cortex (PFC) responded more to the no-go than the infrequent- or frequent-go conditions (Figure 2; sighted: no-go vs. frequent-go t(18)=5.58, p<0.001; no-go vs infrequent-go t(18)=3.15, p=0.006; blind: no-go vs. frequent-go t(18)=4.28, p<0.001; no-go vs infrequent-go t(18)=3.44, p=0.003). We also observed higher responses to the infrequent-go than the frequent-go condition in the rPFC (sighted: t(18)=5.82, p<0.001; blind t(18)=2.15, p=0.045). Responses in the rPFC did not differ between groups (group x condition ANOVA: main effect of condition F(2,72)=33.91, p<0.001; main effect of group F(1,36)=0.01, p>0.5; group x condition interaction F(2,72)=0.39, p>0.5).

**Figure 1.**
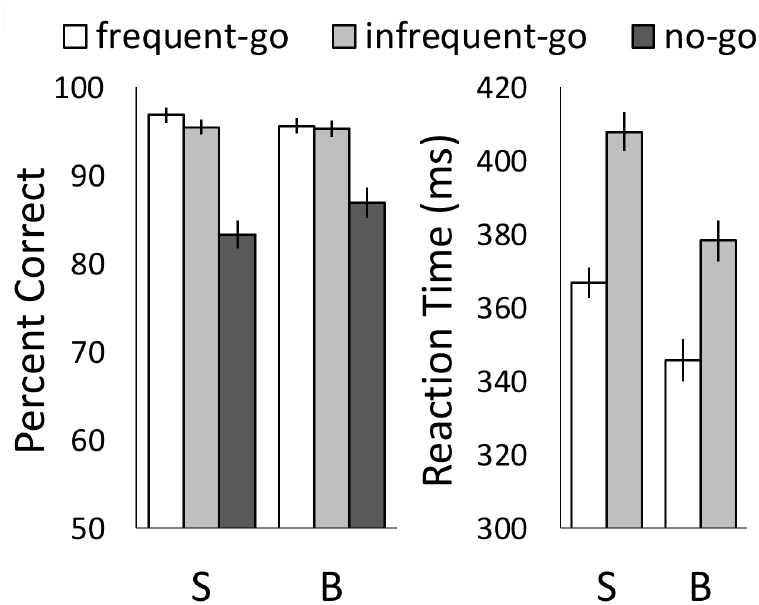
Behavioral performance. Percent correct and response times for sighted (S) and blind (B) participants. Error bars indicate the within-subjects standard error of the mean.

**Figure 2.**
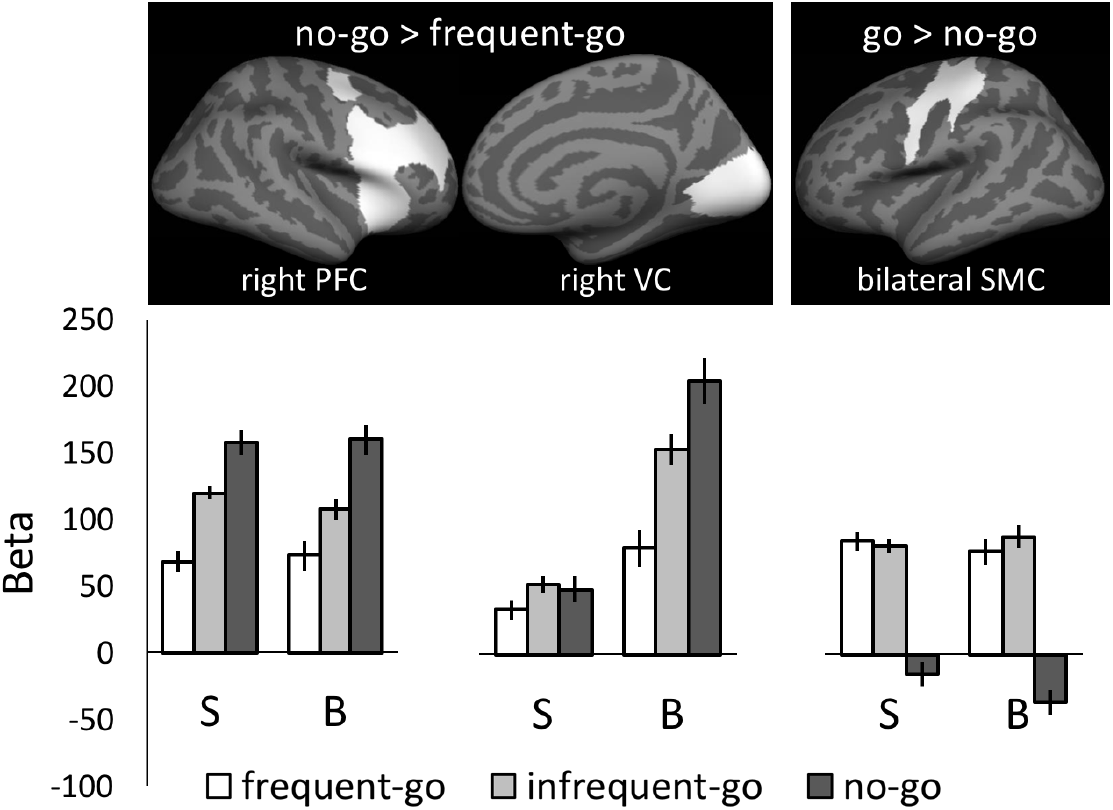
Beta values for task conditions within sighted (S) and congenitally blind (B) participants’ prefrontal cortex (PFC), medial visual cortex (VC), and sensorimotor cortex (SMC). Error bars indicate the within-subjects SEM.

In blind individuals, the right retinotopic visual cortex (VC; i.e. V1-V3) showed a functional profile consistent with the executive function pattern observed in the right PFC: greater response to no-go than frequent-go, a greater response to the infrequent-go than the frequent-go, and a (trending) greater response to no-go than infrequent-go (Figure 2; no-go vs frequent-go t(18)=4.33, p<0.001; infrequent-go vs. frequent-go t(18)=3.75, p=0.001; no-go vs infrequent-go t(18)=1.99, p=0.06). By contrast, we did not observe this profile of response in the visual cortices of sighted blindfolded controls (Figure 2; no-go vs frequent-go t(18)=0.96, p = 0.35; no-go vs infrequent-go t(18)=0.24, p>0.5; infrequent-go vs. frequent-go t(18)=1.77, p=0.09; condition x group ANOVA: main effect of condition F(2,72)=12.30, p < 0.001, main effect of group F(1,36)=7.01, p=0.01, group x condition interaction F(2,72)=7.31, p=0.001).

Within the congenitally blind group’s “visual” cortices, the executive function profile was more pronounced in the right hemisphere than left hemisphere (hemi x condition ANOVA: main effect of condition, F(2,36)=9.37, p = 0.001; main effect of hemi F(1,18)=2.34, p=0.14, hemi x condition interaction F(2,36)=3.59, p = 0.04). Likewise, a hemispheric difference with respect to condition was also present in the PFC (blind group only, hemi x condition ANOVA: main effect of condition, F(2,36)=12.57, p<0.001; main effect of hemi F(1,18)=4.97, p = 0.04, hemi x condition interaction F(2,36)=4.75, p=0.015). Within the blind group, prefrontal and “visual” cortices behaved similarly. There was no difference between the PFC and the VC with respect to condition and/or hemisphere (ROI x hemi x condition ANOVA: main effect of ROI F(1,18)=2.66, p=0.12; ROI x condition interaction F(2,36)=2.53, p=0.09; ROI x hemi interaction F(1,18)=0.50, p=0.49; ROI x hemi x condition F(2,36)=0.09, p>0.5).

### Primary sensory-motor, but not “visual” cortices, of blind individuals respond to sensorimotor demands (individual-subject functional ROI analysis)

In the bilateral sensory-motor cortices (SMC), we observed higher activity for both of the go conditions than the no-go in both the sighted (Figure 2; frequent-go vs no-go t(18)=6.20, p<0.001; infrequent-go vs no-go t(18)=7.47, p<0.001; frequent-go vs. infrequent-go t(18)=0.42, p=0.68) and the blind group (frequent-go vs no-go t(18)=6.61, p < 0.001; infrequent-go vs. no-go t(18)=8.29, p<0.001; frequent-go vs. infrequent-go t(18)=0.68, p=0.51). This response profile is consistent with SMC involvement in execution of the button press and associated tactile feedback. There was no difference between go conditions in the SMC for either group (sighted: t(18)= 0.42, p > 0.5; blind t(18)=0.68, p>0.5). The SMC profile was similar in blind and sighted individuals (group x condition ANOVA: main effect of condition F(2,72)=73.64, p<0.001; main effect of group F(1,36)=0.05, p>0.5; group x condition interaction F(2,72)=0.87, p=0.42).

In contrast to the SMC, we failed to observe a sensorimotor related effect in the right visual cortices of blind individuals even when we searched specifically for vertices that preferred trials with a button press (frequent and infrequent go) to no-go trials (Figure S1, frequent-go vs no-go t(18)=-0.96, p=0.35; infrequent-go vs no-go t(18)=1.40, p=0.18; frequent-go vs. infrequent-go t(18)=2.33 p=0.03). Interestingly, in the sighted group there was a trend towards higher responses to the two button press conditions (Figure S1, frequent-go vs no-go t(18)=2.07, p=0.053; infrequent-go vs no-go t(18)=2.03, p=0.057; frequent-go vs. infrequent-go t(18)=0.13 p>0.5).

### Whole-brain Analysis

Consistent with the ROI analysis and with previous findings, the no-go > frequent-go contrast revealed robust responses in right-lateralized, prefrontal and parietal executive function networks of both blind and sighted groups (Figure 3A). Greater activity for no-go than frequent-go was observed along the precentral sulcus, inferior frontal sulcus, inferior frontal junction (IFJ), and intraparietal sulcus (IPS), as well as the superior temporal sulcus and gyrus (STG/STS; Figure 3A & Table S1). Additionally, we observed greater activity for the no-go condition in the supplementary motor area/anterior cingulate cortex (SMA/ACC) of the blind group and in the posterior precuneus of the sighted group. In the blind and sighted groups, responses to no-go > frequent-go, were observed bilaterally but were stronger on the right (Figure S2).

**Figure 3.**
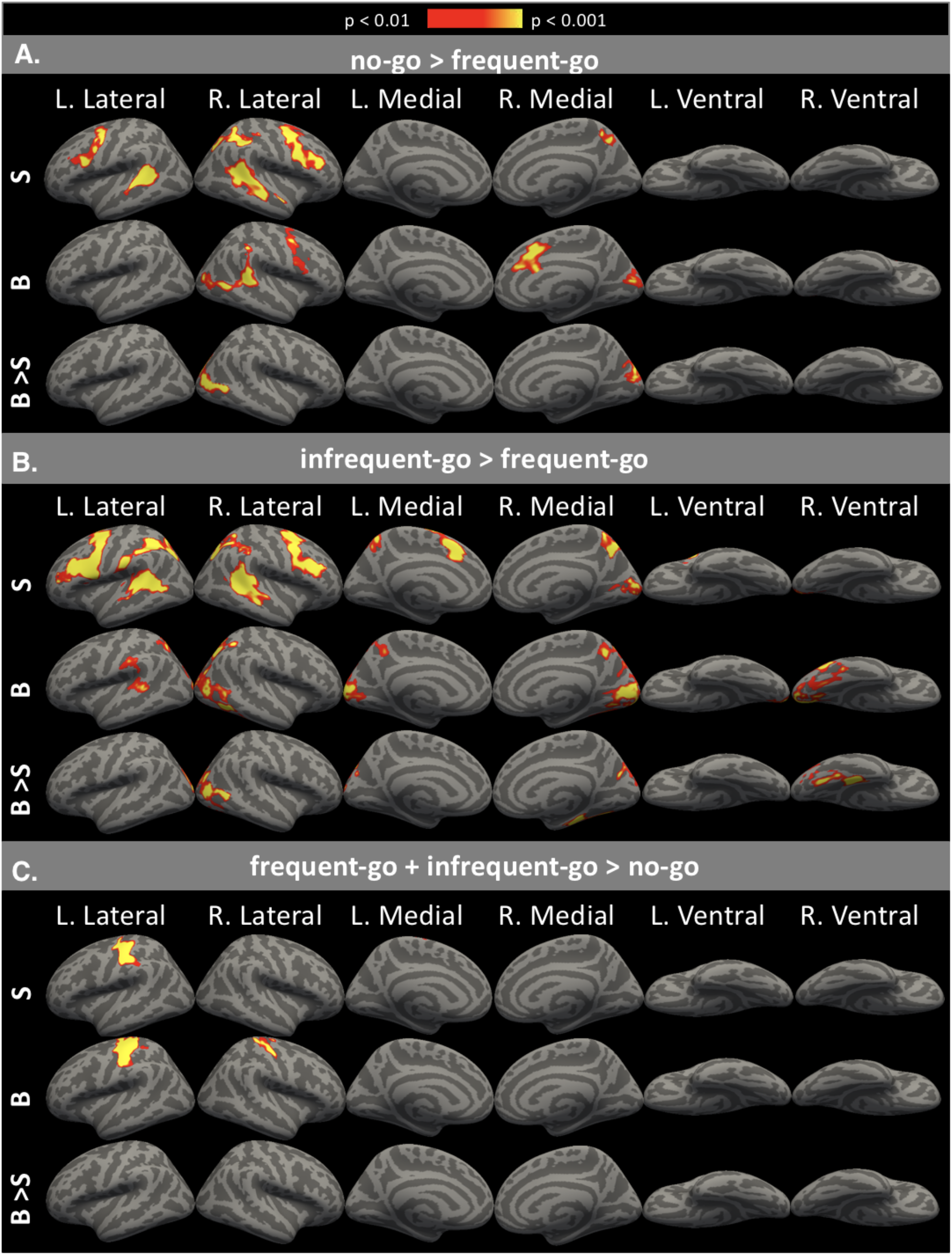
Whole brain contrasts for sighted (S), blind (B), and blind > sighted (B > S). Areas shown are p < 0.05 cluster-corrected p-values, with intensity representing uncorrected vertex-wise probability.

Similar to no-go, infrequent-go also elicited greater activity in executive function regions, relative to frequent-go in both the sighted and blind groups (Figure 3B & Table S1). In the sighted group, the same fronto-parietal and temporal areas that responded more to no-go than frequent-go also responded more to infrequent-go than frequent-go. Notably, fronto-parietal networks were recruited more bilaterally for infrequent-go than for no-go. In the blind group, infrequent-go > frequent-go activity was observed in parts of the IPS, STS, and posterior precuneus. Fronto-parietal responses to executive demands were somewhat less extensive in the blind relative to the sighted group.

In the blind but not sighted group, retinotopic “visual” cortices responded more to the no-go than to the frequent-go condition (Figure 3A & Table S1). Activity in the occipital cortices of the blind group mirrored that of fronto-parietal cortices in right-hemisphere dominance. Occipital cortex activity, in the blind group, peaked in the cuneus and the lateral middle occipital gyrus. Comparing blind and sighted groups directly, we observed greater activity in the congenitally blind group, for no-go relative to frequent-go, on the lateral and medial surface of the right occipital cortex (Figure 3A & Table S1).

The infrequent > frequent go contrast also revealed activity in occipital cortices, but this time in both the blind and sighted groups. In the blind group, the anatomical distribution of the infrequent > frequent go response overlapped with the “no-go” response in the cuneus and lateral medial occipital gyrus but also extended into the right fusiform gyrus and the calcarine sulcus bilaterally. As in prefrontal cortices, occipital cortices exhibited reduced right-lateralization for the infrequent-go > frequent-go contrast, relative to the no-go > frequent-go contrast. In the sighted group, infrequent > frequent go activity was anatomically constrained to the posterior calcarine sulcus—i.e. the functional location of foveal V1—and bilateral (Figure S2). When blind and sighted groups were compared to each other directly, there was greater activity in the blind group in lateral occipital cortices as well as the medial fusiform (Figure 3B, Table S1).

Primary sensory and motor cortices, but not occipital cortices, responded to sensorimotor demands of frequent-go and infrequent-go. For both blind and sighted groups, left-hemisphere primary sensory and primary motor cortices were more active for both go conditions than for the no-go condition (Figure 3C & Table S1). For the blind group, greater activity for the go conditions was also observed in the right primary sensory and primary motor cortices, consistent with fact that more blind individuals using their left hand to respond (see Methods). Consistent with the ROI analysis, no “visual” cortex activity was observed in the blind group for frequent- and infrequent-go conditions relative to the no-go condition. Moreover, a direct comparison between blind and sighted groups revealed no interaction of group by condition.

### Resting State Functional Connectivity

We used resting state data to examine functional connectivity of executive-load responsive “visual” cortex among blind individuals. An executive-function responsive visual ROI (OC-EF) was defined based on the blind > sighted x no-go > frequent-go contrast (see Methods for details). For both blind and sighted participants, we assessed OC-EF connectivity to two primary sensory regions—A1 and S1/M1— and to two prefrontal regions—one responsive to executive-function, PFC-EF, and one responsive to language, PFC-LG (Figure 4A).

**Figure 4.**
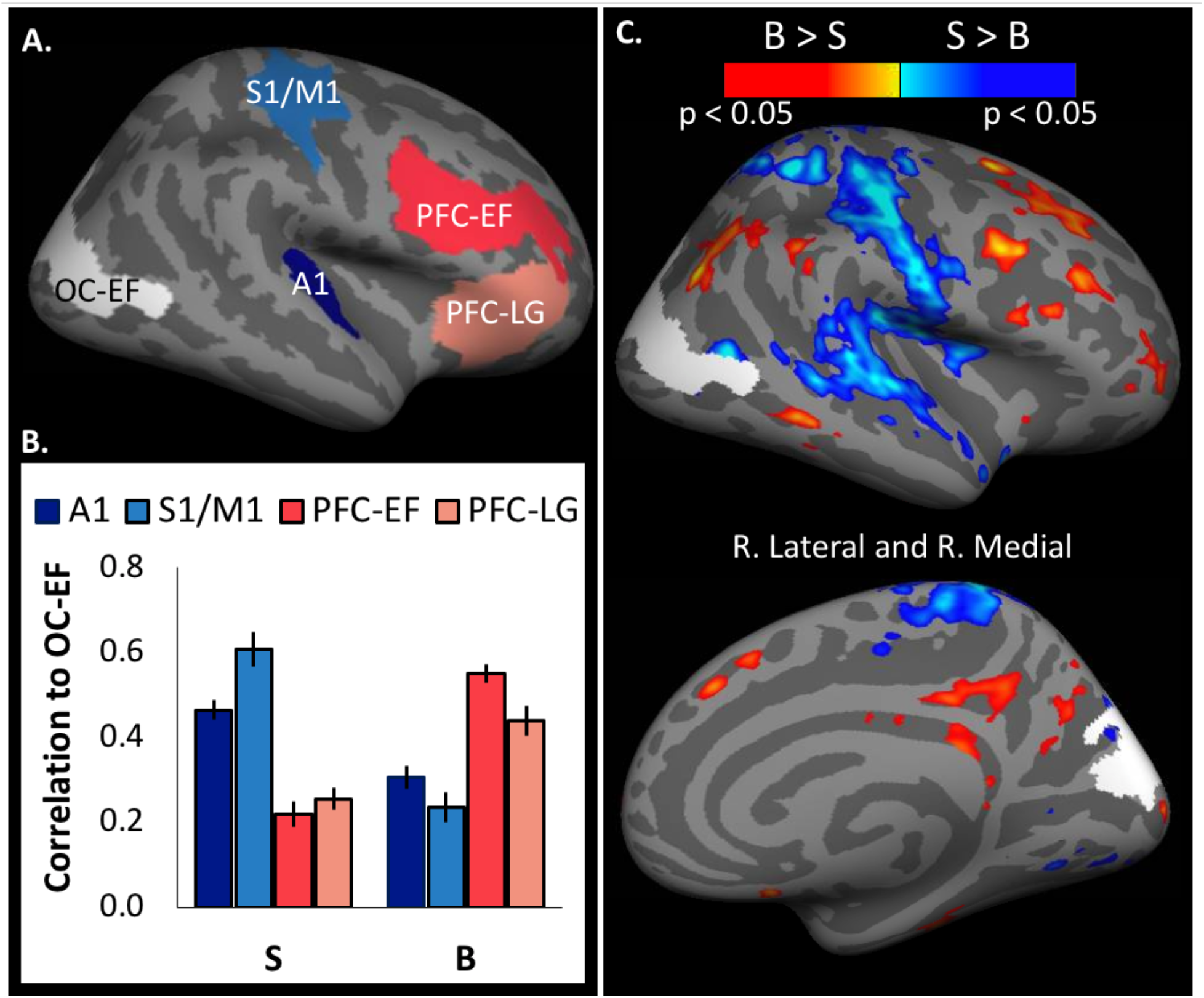
Functional connectivity of executive-function responsive occipital cortex (OC-EF) to sensory/motor and prefrontal regions in sighted and blind. *A*. Regions of interest for resting state analyses: OC-EF, primary auditory cortex (A1), primary sensorimotor cortices (S1/M1), executive-function responsive (PFC-EF), and language responsive (PFC-LG). *B*. Fisher-transformed correlation coefficients between OC-EF and non-visual ROIs. Error bars indicate the within-subjects SEM. *C*. Between-group differences in connectivity to executive-function responsive occipital cortex (OC-EF, in white). FDR-corrected contrasts for blind > sighted (in red) and sighted > blind (in blue).

An ANOVA comparing the connectivity of executive-function responsive visual cortex (OC-EF) to executive-function responsive prefrontal, language-responsive prefrontal, sensory-motor and primary auditory ROIs across groups revealed a significant group by ROI interaction (ROI (PFC-EF, PFC-LG, A1, S1/M1) x group ANOVA: main effect of ROI F(3,144)=1.48, p=0.22; main effect of group F(1,48)=0.01, p>0.5; group x ROI interaction F(3,144)=41.63, p<0.001). An ANOVA within sighted individuals revealed that the executive function responsive visual cortex (OC-EF) was more correlated to non-visual primary sensory areas (A1 and M1/S1) than to either of the executive function or language-responsive prefrontal ROIs (mean of PC-EF and PC-LG < mean of A1 and M1/S1: F(1,24)=42.98, p<0.001). Conversely, in blind individuals, OC-EF was more correlated to prefrontal than to primary sensory regions (mean of PC-EF and PC-LG > mean of A1 and M1/S1: F(1,24)=21.99, p<0.001). Finally, among the prefrontal cortex ROIs, OC-EF of blind but not sighted individuals was preferentially correlated to executive function-responsive prefrontal cortex than language-responsive prefrontal cortex (blind PC-EF vs. PC-LG t(24)=3.47, p=0.002, sighted PC-EF vs. PC-LG t(24)=-1.17, p=0.25; group x ROI (PFC-EF vs. PFC-LG) interaction F(1,48)=10.86, p=0.002).

Next, we compared OC-EF functional connectivity between groups throughout the whole-brain. Relative to sighted individuals, blind individuals had increased OC-EF connectivity to fronto-parietal regions and decreased OC-EF connectivity to primary sensory areas (Figure 4C). Moreover, for blind individuals, the set of regions that preferentially increased their correlation to OC-EF at rest was equivalent to the set of regions that exhibited an executive-function response profile during the go/no-go task. Areas that were more functionally connected to the OC-EF for blind individuals included the precentral sulcus, superior and inferior frontal sulcus, inferior frontal junction (IFJ), intraparietal sulcus (IPS), middle temporal gyrus (MTG), precuneus, and the anterior and posterior cingulate cortex (ACC and PCC). In contrast, the pre-to-post central gyrus, transverse temporal gyrus, and ventral superior temporal gyrus (STG) were more functionally connected to the OC-EF in sighted, than in blind, individuals

## Discussion

Two key findings support the hypothesis that a right-lateralized subset of “visual” cortex is incorporated into a right-lateralized fronto-parietal non-verbal executive function network in congenital blindness. First, in blind individuals, a right-lateralized sub-network within “visual” cortices is sensitive to executive demands during a non-verbal, non-spatial, go/no-go task. The occipital cortices of congenitally blind (but not sighted) adults were most active during no-go trials, i.e. when withholding a button press. Amongst the go trials, responses in the “visual” cortices of the blind group were larger for the infrequent-go condition than for the frequent-go condition. This response-profile (no-go > infrequent-go > frequent-go) mirrored that observed in the fronto-parietal executive function network of both blind and sighted groups. Second, in blindness executive-function responsive “visual” cortex becomes functionally coupled with prefrontal executive function areas, even at rest.

### Visual cortices of congenitally blind individuals are sensitive to executive demands in a nonverbal and non-spatial task

We find that in blindness, regions of the “visual” cortices are sensitive to non-verbal executive demands. These responses are functionally and anatomically distinct from several previously documented cross-modal effects. As noted in the introduction, “visual” cortices of blind individuals show sensitivity to linguistic information and to mathematical difficulty (Bedny et al., 2011; Kanjlia et al., 2016; Lane et al., 2015; Röder et al., 2002). However, the present stimuli were not linguistic or mathematical. Furthermore, previously observed “visual” cortex responses to language are on-average left-lateralized and responses to math were observed bilaterally (Bedny et al., 2011; Kanjlia et al., 2016; Lane et al., 2015). In contrast, executive-function responses observed in the current study are lateralized to the right-hemisphere. Furthermore, while mathematical responses were limited to the posterior “visual” cortices, executive function responses were observed laterally and medially, as well as posteriorly. Different cortical locations for executive function responses, compared to language or math responses, suggests functionally distinct repurposing in the “visual” cortices. Future work should test for functionally-selective sub-regions within the “visual” cortices of blind individuals, rather than at the group level.

Low-level sensorimotor demands are also unlikely to explain executive function responses in the “visual” cortices observed in the current study. First, the condition that elicited the most activity (i.e. no-go) had the highest executive demand but the lowest tactile feedback and motor planning demand. Second, we failed to find a response profile within the “visual” cortices of blind individuals akin to the response of sensorimotor cortices. Previous studies with blind participants have also failed to find “visual” cortex activity during low-level sensorimotor tasks, such as passive vibro-tactile stimulation, tactile sweeping of non-sense Braille without discrimination, and finger tapping, despite task recruitment of sensorimotor cortices (Gizewski, Gasser, de Greiff, Boehm, & Forsting, 2003; Sadato et al., 1996). Similarly, passively presented visual and tactile stimuli failed to elicit “auditory” cortex activity in a congenitally deaf participant, despite recruitment of primary visual and sensorimotor cortices, respectively (Hickok et al., 1997). Together with these prior results, our findings suggest that “visual” cortices, are not likely to be repurposed for primary sensory-motor functions in blindness.

Finally, responses to executive demands observed in the current study are unlikely to be related to spatial processing. As noted in the introduction, previous studies have observed “visual” cortex responses during tasks that require localization— e.g. localization of sounds in space and discrimination of tactile patterns (Collignon et al., 2011; Gougoux et al., 2005; Merabet et al., 2004; Sadato et al., 1996; Sathian & Stilla, 2010; Wanet-Defalque et al., 1988). By contrast, in the current experiment, auditory stimuli were not situated in space and the task did not require localization nor did the task involve fine-grained tactile discrimination. The present findings thus demonstrate that executive-demands influence visual cortex activity independent of spatial processing. We hypothesize that, in blindness, spatial processing engages different subsets of “visual” cortices as compared to executive, linguistic, and numerical processes.

In future work, it will be important to determine the precise nature of the executive operations that drive activity in deafferented “visual” cortices. Executive processes include a diverse set of operations, such as response selection, response inhibition, shifting attention, and saliency orienting. According to some views, these subtypes of executive control are dissociable within fronto-parietal cortices (Aron et al., 2004; Brass, Derrfuss, Forstmann, & Cramon, 2005; Chikazoe, 2010; Chikazoe et al., 2008; Corbetta & Shulman, 2002; Goghari & MacDonald, 2009; Levy & Wagner, 2011; Mostofsky et al., 2003; Nagahama et al., 2001; Nee, Wager, & Jonides, 2007; Rubia, Smith, Brammer, & Taylor, 2003; Xu et al., 2017). In the current study, “visual” cortices responded to both stimulus infrequency (infrequent-go and no-go) and response infrequency/inhibition (no-go). One possibility is that “visual” cortices are specifically sensitive to response inhibition, and that intermediate activity for the infrequent-go is reflective of participants “tapping the brakes” on their go response (as in a “continue” trial; (Aron et al., 2014)). Alternatively, the observed executive functions responses may reflect a response selection process that scales according to novelty of the stimulus-response mapping (i.e. frequent-go has both a habitual stimulus and a habitual response, infrequent-go has a novel stimulus but a habitual response, and no-go has both a novel stimulus and a novel response; (Chikazoe et al., 2008)). Whether the “visual” cortex contain dissociable response selection and response inhibition processes and whether it is sensitive to other types of executive processes are interesting avenues for future research.

A further key question to be addressed in future work concerns the behavioral relevance of “visual” cortices to executive function and other higher-cognitive processes. “Visual” cortices may be repurposed to perform computations typical of fronto-parietal cortices, but can they influence behavior? There is evidence that “visual” cortices of blind individuals are relevant to some higher-cognitive tasks, such as Braille reading and verb generation (Amedi, Floel, Knecht, Zohary, & Cohen, 2004; Amedi, Raz, Pianka, Malach, & Zohary, 2003; L. G. Cohen, Celnik, Pascual-Leone, & Corwell, 1997). For example, TMS to “visual” cortices causes blind individuals to make semantic errors when asked to generate a verb to an auditory presented noun (i.e. “kick” for “ball”; (Amedi et al., 2004)). In the current study, the blind group did not significantly outperform the sighted group on the go/no-go task. One possibility is that “visual” cortices do not confer a response selection benefit. Alternatively, blind individuals could show enhanced performance in a more demanding task. Previous studies have found that blind individuals outperform the sighted in tasks of verbal working memory and divided attention (Amedi et al., 2003; Collignon, Renier, Bruyer, Tranduy, & Veraart, 2006; Dormal, Crollen, Baumans, Lepore, & Collignon, 2016; Hull & Mason, 1995; Roder, Rösler, & Neville, 2001; Rokem & Ahissar, 2009; Swanson & Luxenberg, 2009; Tillman & Bashaw, 1968; Withagen, Kappers, Vervloed, Knoors, & Verhoeven, 2013). Whether “visual” cortex are functionally relevant to executive processes and whether this functional relevance underlies previously observed behavioral advantages remains to be tested in future research.

Interestingly, in the current study we also observed responses to non-visual stimuli in the visual cortices of blindfolded, sighted adults. Importantly, these responses were functionally and anatomically distinct from those observed in the “visual” cortices of blind individuals. While the “visual” cortices of the blind group showed a graded executive demand response (with the highest response to no-go and the lowest response to frequent-go), the visual cortices of the sighted group showed selective high responses to the infrequent-go condition. Moreover, while “visual” cortex responses in the blind group were predominantly right-lateralized and extended along the medial, lateral, and ventral surface, visual cortex responses in the sighted group were bilateral and strictly localized to the calcarine sulcus (V1).

The cognitive role of visual cortex responses to non-visual stimuli in sighted individuals is not known. Some prior studies have also observed responses to non-visual stimuli in visual cortex of sighted subjects, for example, in participants who are trained to associate a visual flash with an auditory noise, visual cortex activity is observed during subsequent presentation of the noise alone (Driver & Noesselt, 2008; James et al., 2002; Macaluso, Frith, & Driver, 2000; Merabet et al., 2008; 2004; Sathian, Zangaladze, Hoffman, & Grafton, 1997; Zangaladze, Epstein, Grafton, & Sathian, 1999; Zangenehpour & Zatorre, 2010). One possibility is that the visual cortex responses in sighted subjects observed in the current study reflect automatic orienting. It has been hypothesized that unexpected auditory stimuli elicit a “reflexive overt orienting response” towards the location of visual space where the stimulus is most likely to occur (Azevedo, Ortiz-Rios, Li, Logothetis, & Keliris, 2015). According to this account, in the absence of further spatial information, there is automatic orienting to the center of the visual field and pre-activation of foveal V1 specifically in cases of planning a motor action and when the stimulus response mapping is not highly overlearned (i.e., as in infrequent-go). At present these interpretations are speculative and will require testing in future research. However, such effects are consistent with the idea that there are routes for non-visual information to reach “visual” cortex in blind and sighted alike and these routes are modified by absence or lack of visual experience.

### Insights into the relationship of connectivity and function from “visual” cortex plasticity in blindness

Further support for the idea that, in blindness, parts of right “visual” cortices are incorporated into fronto-parietal executive function networks comes from resting state data. The executive-function responsive “visual” cortex of blind individuals was coupled with executive-load responsive prefrontal cortices. Specifically, blind and sighted groups displayed different profiles of functional connectivity for the occipital cortex area in which executive-function responses were observed in blindness. In the sighted group, executive-function responsive visual cortex was more correlated with non-visual sensory-motor areas (A1 and M1/S1) than with prefrontal cortices. Conversely, in the blind group, executive-function responsive “visual” cortex was more correlated with prefrontal cortices than with non-visual sensory-motor areas. This change in connectivity was driven both by a reduction in resting state correlations with A1 and S1/M1 as well as an increase in correlations with prefrontal cortices in blindness. This result is consistent with prior studies, which have also found reduced connectivity of “visual” cortices, in blindness, to A1 and to sensory-motor cortices (Burton, Snyder, & Raichle, 2014; Y. Liu et al., 2007; Wang et al., 2013; Yu et al., 2008). By contrast, resting-state synchrony between “visual” cortices and frontal-parietal cortices is increased in blindness (Bedny et al., 2011; Bedny, Konkle, Pelphrey, Saxe, & Pascual-Leone, 2010; Burton et al., 2014; Deen, Saxe, & Bedny, 2015; L. Liu et al., 2017; Y. Liu et al., 2007; Wang et al., 2013).

Importantly, among prefrontal regions, executive-load responsive “visual” cortex was more correlated with executive-function responsive prefrontal cortex, than language responsive prefrontal cortex, and this effect was specific to the blind group. This result supports the hypothesis that the executive load responsive “visual” areas identified in the present study are functionally distinct from previously identified language-responsive visual areas. Analogously, previous studies have found that prefrontal language areas are more synchronized to the parts of “visual” cortices that respond to language than to the parts of “visual” cortices that respond to math (Kanjlia et al., 2016). These results demonstrate that resting-state connectivity and functional specialization within “visual” cortex go hand in hand in blindness.

Together, the available resting-state and task-based findings from blindness support the hypothesis that anatomical connectivity plays a major role in driving cortical function. The finding that occipital cortices of blind individuals take on fronto-parietal functions is consistent with the observation that, in sighted and blind individuals alike, fronto-parietal networks constitute a main source of anatomical afferent connections to the visual system (Bressler et al., 2008; Gilbert & Li, 2013). Since there is no evidence of large-scale additional anatomical tracts in blind relative to sighted individuals, these functional changes are likely to result from long-range connectivity between fronto-parietal networks and visual cortices that are present in both blind and sighted groups (Shimony et al., 2005; Shu, Li, Li, Yu, & Jiang, 2009a; Shu et al., 2009b). We hypothesize that the functional reorganization observed in blindness is mediated by local synaptic changes that alter the efficacy of top-down anatomical inputs from higher-cognitive regions.

Further support for the idea that long-range anatomical connectivity constrains the function of cortex comes from the localization of different functions within the visual cortices of blind individuals. Across multiple examples of plasticity, “visual” cortex functions in blind individuals are co-lateralized with the non-visual cortices that classically implement such functions. In the current study, executive function activity (during the go/no-go task) was right lateralized in both fronto-parietal and “visual” cortices. By contrast, language responses in the “visual” cortices are on average more pronounced in the left hemisphere, in keeping with left lateralization of language in frontotemporal cortices (Bedny et al., 2011; Lane et al., 2015; Röder et al., 2002). Moreover, in blind individuals with right-lateralized language processing in the fronto-temporal cortices, language responses in the “visual” cortices are also right-lateralized (Lane et al., 2017). Because anatomical connectivity is stronger within, than across, hemispheres, co-lateralization of blind “visual” cortices is consistent with the hypothesis that plasticity is constrained by pre-existing anatomical connections to the occipital cortices.

Support for anatomical connectivity-based constraints on function also comes from studies outside of blindness (Dehaene & Cohen, 2007; Johnson, 2000; Mahon & Caramazza, 2011; O’Leary, 1989). For example, anatomical connectivity predicts which region of the ventral object-recognition stream will become the “visual word form area” (VWFA) (Z. M. Saygin et al., 2016). Relative to other parts of the ventral stream, this cortical location has strong reciprocal anatomical connectivity with fronto-temporal language networks, even prior to onset of literacy (Dehaene, Cohen, Morais, & Kolinsky, 2015). Such results are consistent with findings from studies of blindness. Together these studies support the view that anatomical connectivity plays a major role in shaping cortical function.

Studies of blindness demonstrate that anatomical connectivity constraints on cortical function are not inconsistent with the possibility of large-scale functional flexibility. Anatomical connectivity constrains the range of cognitive functions a cortical area can assume by regulating its input. Different regions of cortex receive inputs from distinct sensory modalities and cognitive domains and this specialized input has powerful effects on cortical function. However, because the intrinsic microcircuitry of human cortex is cognitively flexible at birth, experience can drastically alter the consequences of these anatomical biases. The same anatomical connectivity pattern that mediates communication between vision and higher-order cognition in those who are sighted enables the incorporation of occipital cortices into higher cognitive networks in blindness.

## Materials and Methods

### Participants

19 congenitally blind and 19 sighted controls (blind: 13 females; 12 right-handed, 3 ambidextrous; age: mean=45.3, s.d.=17.43; years of education: mean=17.00, s.d.=2.73; sighted: 14 females; 18 right-handed; age: mean=41.71, s.d.=14.74; years of education: mean=17.97, s.d.=3.68) contributed task-based data. Blind and sighted participants were matched on average age (t(36)=0.50, p=0.69) and education level (t(36)=0.36, p=0.93).

All but one sighted participant from the task-based go/no-go experiment contributed resting state data. Resting state data from an additional 6 blind and 7 sighted participants were included, resulting in the following group-wise demographics (blind: N=25; 18 females; age: mean = 46.63, s.d.=16.9; sighted: N=25; 15 females; age: mean = 43.16, s.d.=12.26; blind vs. sighted age, t(48)=0.83, p=0.41). During the resting state scan, participants were instructed to relax but remain awake.

All blind participants self-reported minimal-to-no light perception since birth, i.e. having never been able to distinguish colors, shapes, or motion. (One blind participant was born with no light perception, but reported some functional vision in one eye between 5 and 11 years of age, following several corrective surgeries. This participant’s data was no different from the remaining blind group and is included in the sample.) Blind and sighted participants had no known neurological disorders, head injuries, or brain damage. For all blind participants, the causes of blindness excluded pathology posterior to the optic chiasm (see Table 1 for details). All participants gave written consent under a protocol approved by the Institutional Review Board of Johns Hopkins University. All participants wore light exclusion blindfolds for the duration of the scan to equate light conditions across participants.

**Table 1.** Per cause of blindness, total number of participants (No.) and number with light perception (LP No.). Amounts outside of parentheses are for participants in task-based go/no-go experiment. Amounts within parentheses are for additional participants included in resting-state analyses.

| **Blindness Etiology** | **No.** | **LP**<br>**No.** |
| --- | --- | --- |
| Leber Congenital Amaurosis | 5 (+2) | 4 (+2) |
| Retinopathy of Prematurity | 8 (+4) | 4 |
| Optic Nerve Hypoplasia | 2 | 0 |
| Retinitis Pigmentosa | 1 | 1 |
| Glaucoma | 1 | 0 |
| Unknown | 2 | 1 |

### fMRI data acquisition

MRI structural and functional data of the whole brain were collected on a 3 Tesla Phillips scanner. T1-weighted structural images were collected in 150 axial slices with 1 mm isotropic voxels using a magnetisation-prepared rapid gradient-echo (MP-RAGE). T2*-weighted functional images were collected in 36 axial slices with 2.4 × 2.4 × 3 mm voxels and a 2 second TR. We acquired 3 runs of task-based fMRI data per subject and between 1 and 4 runs of resting state data (mean number of runs: sighted = 1.28, blind =2.08; t(48) = 3.78, p < 0.001).

Acquisition parameters were identical for resting and task-based data.

### Behavioral Task

Participants heard complex non-verbal sounds (450 MS with 450 MS ISI), each representing 1 of 3 conditions: frequent-go (50% trials), no-go (25%), and infrequent-go (25%) and were asked to make speeded button presses in response to the go sounds and to withhold responding to the no-go sounds. Each run was comprised of 400 trials and 4 20-second rest periods, spaced equidistantly, for a total time of 7.67 minutes per run. The full experiment consisted of three runs. Presentation order was constrained so that each infrequent condition – i.e. infrequent-go and frequent-go – could not occur on more than 3 consecutive trials. Feedback on performance accuracy was given after every run. To avoid making participants explicitly aware of the frequency manipulation, the frequent-go and infrequent-go conditions were referred to as “go 1” and “go 2,” respectively. Prior to taking part in the main experiment, participants performed 400 practice trials (100 inside the scanner) with auditory feedback after each trial.

The 3 stimulus sounds were chosen to be easily and immediately discerned from each other. All 3 sounds differed from each other at the sound onset and remained relatively homogenous throughout the sound duration. To discourage chunking of sounds across conditions, sounds were selected to be equally dissimilar (Amazon Mechanical Turk pre-experiment pilot testing revealed no one sound as the “odd one out”; chi-sqd(2)=1.66, N=49; p > 0.5). Assignment of sounds to conditions was counterbalanced across participants and matched across blind and sighted groups.

Auditory stimuli were presented over Sensimetrics MRI-compatible earphones (http://www.sens.com/products/model-s14/) at the maximum comfortable volume for each participant. The volume was adjusted for all stimuli or selectively for a specific stimulus (2 sighted, 2 blind) according to participant’s request. Adjustments did not affect the analyzed data, as they were implemented prior to the first functional run. Participants were free to make responses with their preferred hand (right hand for all but 1 sighted and 4 blind participants).

### fMRI task-based data analysis

Data analyses were performed using FSL, Freesurfer, the HCP workbench, and custom software (Dale, Fischl, & Sereno, 1999; Glasser et al., 2013; S. M. Smith et al., 2004). Functional data were motion corrected, slice-time corrected, high pass filtered with a 128 s cutoff, pre-whitened, and resampled to the cortical surface (discarding subcortical structures and the cerebellum). The data were smoothed with a 12 mm FWHM Gaussian kernel on the surface, which affords better spatial accuracy than comparable smoothing in the volume (Anticevic et al., 2008; Hagler, Saygin, & Sereno, 2006; Jo et al., 2007; Tucholka, Fritsch, Poline, & Thirion, 2012).

For each subject, we defined a GLM to predict BOLD activity according to the following event types, each convolved with the hemodynamic response function: successful frequent-go, successful infrequent-go, successful no-go, failed frequent-go, failed infrequent-go, failed no-go, false starts, and extra button presses. Results report only successful trials. All trial events began at the onset of the auditory stimulus and ended at the offset of the auditory stimulus or the participant’s button press (whichever sensory event ended last).

A covariate of no interest was included to account for motion. Timepoints with framewise-displacement (relative movement) greater than 1.5 mm were excluded by modeling each timepoint as an individual regressor with a value of 1 at the time point and 0 everywhere else (drops per run: blind mean=0.30, s.d.=0.80; sighted mean=0.12, s.d.=0.25; difference between groups t(36)=0.91, p=0.37).

Fixed-effects analyses were used to combine runs within each subject, which were then submitted to a group analyses with subject represented as a random-effect. To control for vertex-wise multiple comparisons, we performed a cluster-wise permutation analysis (Hagler et al., 2006; Nichols & Hayasaka, 2003). Whole-brain maps are first thresholded, and resulting cluster-sizes are then tested against a cluster-size null distribution generated from 5,000 permutations. This approach corrects for multiple comparisons because the each permutation’s null value is determined by the highest cluster size, across the whole brain. Reported whole-brain contrasts were run as one-sided tests, thresholded at p < 0.01 vertexwise, and p < 0.05 cluster-corrected. Because this can eliminate small clusters, we also performed a multiple comparison correction using a false discovery rate (FDR) of 5%, per hemisphere, on one-tailed p-values (Genovese, Lazar, & Nichols, 2002).

We performed individual-subject functional regions of interest (ROIs) by defining three (ROIs) in each participant: 1) a prefrontal (PFC) executive function ROI, 2) a sensory-motor (SMC) ROI and 3) a medial visual cortex (VC). ROIs were defined by selecting responsive vertices for each participant within a group-wise search space using a leave-one run out procedure.

Search spaces were defined as follows. For the executive function PFC ROI and the sensorimotor cortex (SMC) ROI we defined a search based on previous studies that observed response-inhibition and hand-movement activity, respectively, using neurosynth.org (Yarkoni, Poldrack, Nichols, Van Essen, & Wager, 2011). Both volumetric search-spaces were projected to the surface, thresholded at z > 1.65, dilated and eroded at 12 mm (to fill small holes), and restricted to the anatomical area of interest. For the PFC, we confined the functionally derived search space to right lateral prefrontal lobe, anterior to the pre-central gyrus. For the SMC, we confined the functionally derived search space to the left central sulcus and pre- and post-central gyri, as defined by a surface-based atlas (Destrieux, Fischl, Dale, & Halgren, 2010). The visual cortex (VC) search space was defined by combining V1, V2, dorsal V3, and ventral V3 (VP) parcels from the PALS-B12 visuotopic surface-based atlas (Van Essen, 2005). All search spaces were created in the right hemisphere and mirrored to the left hemisphere.

Within each search space, we used a leave-one-run out procedure to define and test subject-specific functional ROIs. The PFC ROI was defined based on the no-go > frequent-go contrast. In the SMC ROI, we selected hypothesized somatosensory and motor responsive vertices using the frequent-go + infrequent-go > no-go contrast. We searched for both PFC-type and SMC-type responses in the VC by defining ROIs based on both contrasts. For all ROIs, we selected the top 20% of vertices that most strongly responded to the contrast of interest in all but one run and extracted signal from the left-out run. Beta-values, for each condition, were obtained by averaging whole-brain GLM Betas across the selected ROI vertices. This procedure was repeated iteratively, leaving out one run at a time, and the resulting Betas were averaged across run iterations, for each subject. ROIs vertices were defined according to a subset of runs and those vertices were assessed on non-overlapping subset of runs. Because ROIs were selected and tested orthogonally for the contrast of interest, participant’s ROIs will only show the contrast of interest if preferential activity replicates across runs (i.e., if not driven by noise).

### Resting State Functional Connectivity Analysis

We used the CONN Toolbox (Whitfield-Gabrieli & Nieto-Castanon, 2012) to compare visual cortex functional connectivity during rest. BOLD data were first smoothed 23 diffusion steps on the surface and registered to MNI-152 standard space. To control for temporal confounds, white matter and cerebrospinal fluid BOLD signals were regressed out and the residual was bandpass filtered (0.008-0.1). Time-courses were first averaged within ROIs and then either correlated to each other (ROI-to-ROI) or to the time-course of each and every vertex (ROI-to-whole-brain).

We defined 1 visual and 4 non-visual cortex group-wise regions of interest. An OC-EF ROI was defined as the largest cluster within the entire occipital cortex that responded more to go than no-go in blind, relative to sighted, participants in the cluster-corrected map. A prefrontal executive-function (PFC-EF) ROI was defined as the area with the largest contiguous all-subject activation for no-go > frequent-go (p < 0.05 FDR-corrected), constrained to the PFC search space. Similarly, a sensorimotor (S1/M1) ROI was selected as the largest contiguous all-subject activation for go > no-go (p < 0.05 FDR-corrected), constrained to the SMC search space. A prefrontal language (PFC-LG) ROI was taken from parcels that have previously been observed to be responsive to linguistic content in sighted subjects (Fedorenko, Hsieh, Nieto-Castanon, Whitfield-Gabrieli, & Kanwisher, 2010). Finally, we selected a primary auditory cortex (A1) ROI as the transverse temporal portion of a gyral based atlas (Desikan et al., 2006; Morosan et al., 2001). All ROI analyses were conducted in the right hemisphere so as to match the hemisphere of the visual cortex OC-EF region. Two ROIs that were originally defined in the left hemisphere (i.e. S1/M1 and PFC-LG) were mirrored to the right-hemisphere. This procedure ensures that any functional connectivity differences amongst ROIs are not driven by differences in connectivity across hemispheres.

## Acknowledgements

We would like to thank the blind and sighted individuals who participated in this research, the blind community and the National Federation of the Blind. Without their support this research would not be possible. This work was supported by grants from the Johns Hopkins Science of Learning Institute, the NIH/NEI (R01 EY027352-01), and a National Science Foundation Graduate Research Fellowship DGE-1232825 (to R.E.L.). We would also like to thank the F. M. Kirby Research Center for Functional Brain Imaging at the Kennedy Krieger Institute.

## Author Contributions

R.E.L. and M.B. came up with the question, designed the experiment, analyzed the data, and wrote the manuscript. R.E.L. collected the data.

---

**Figure S1.**
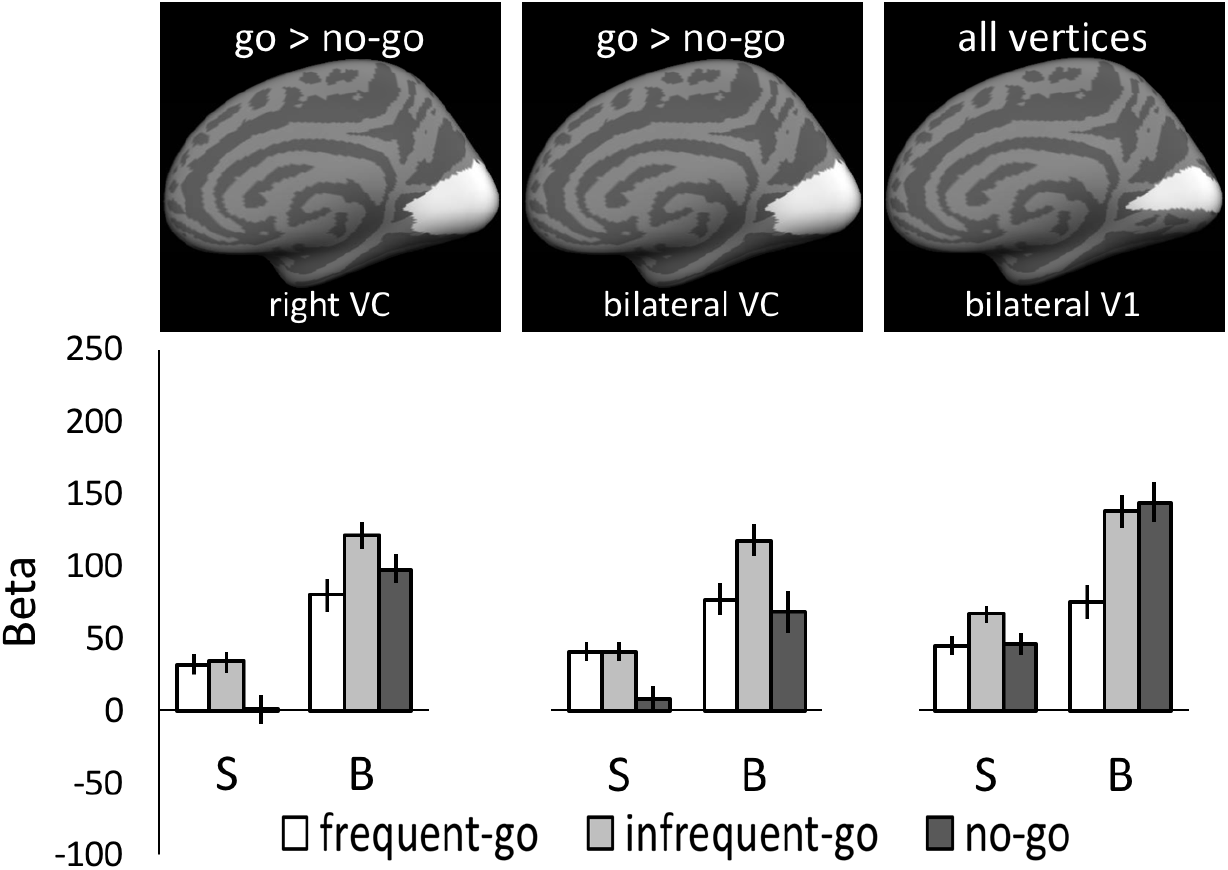
Beta values for task conditions within sighted (S) and congenitally blind (B) participants’ medial visual cortex (VC) and primary visual cortex (V1). Error bars indicate the within-subjects SEM. Right and bilateral VC values are reported from a leave-one out analysis where vertices were chosen based on the contrast frequent-go + infrequent-go > no-go (similar to those reported in the bilateral SMC). Bilateral V1 values were chosen from the entire V1 search-space.

**Figure S2.**
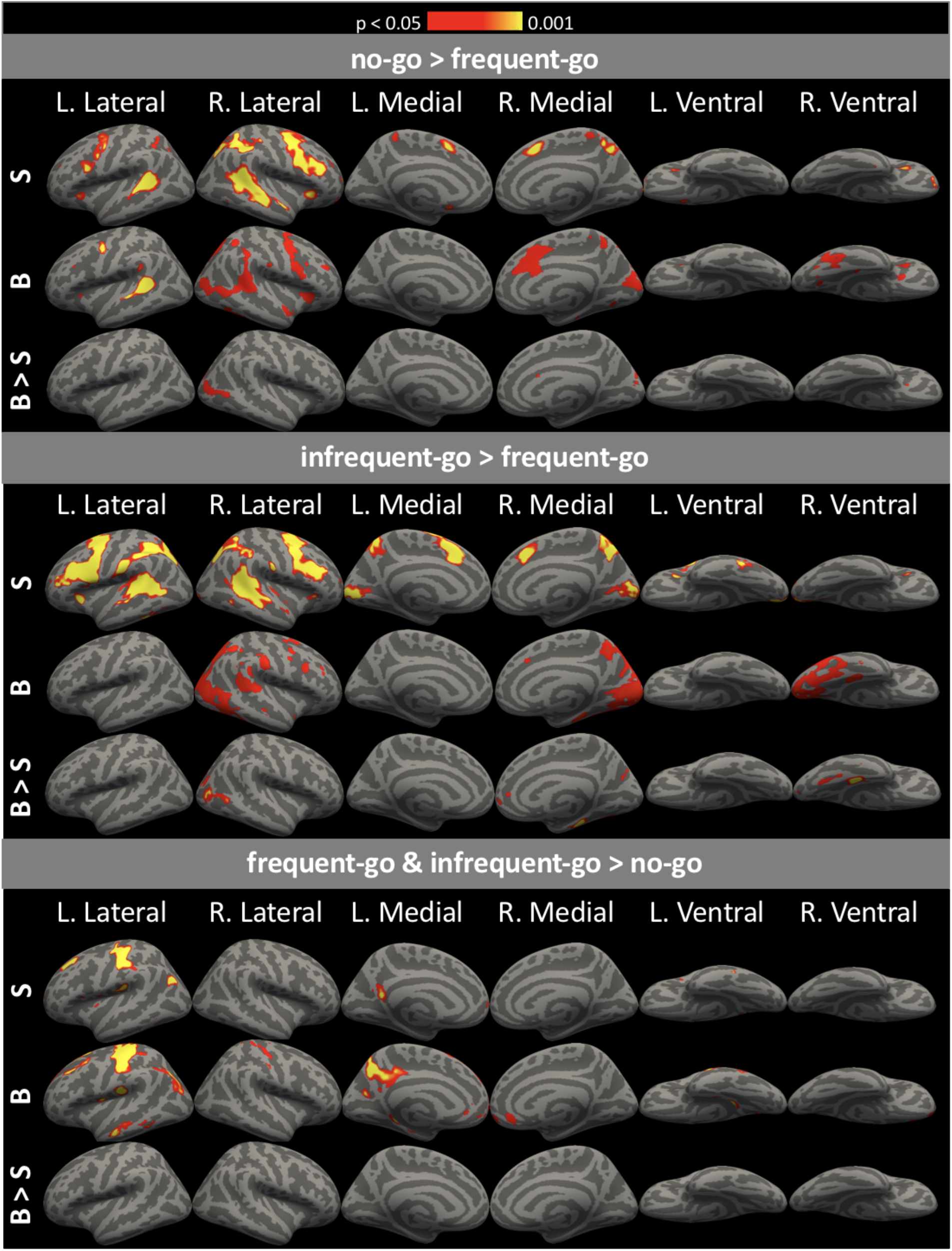
FDR-corrected whole brain contrasts for sighted (S), blind (B), and blind > sighted (B > S). p-values are FDR-adjusted.

**Table S1.**
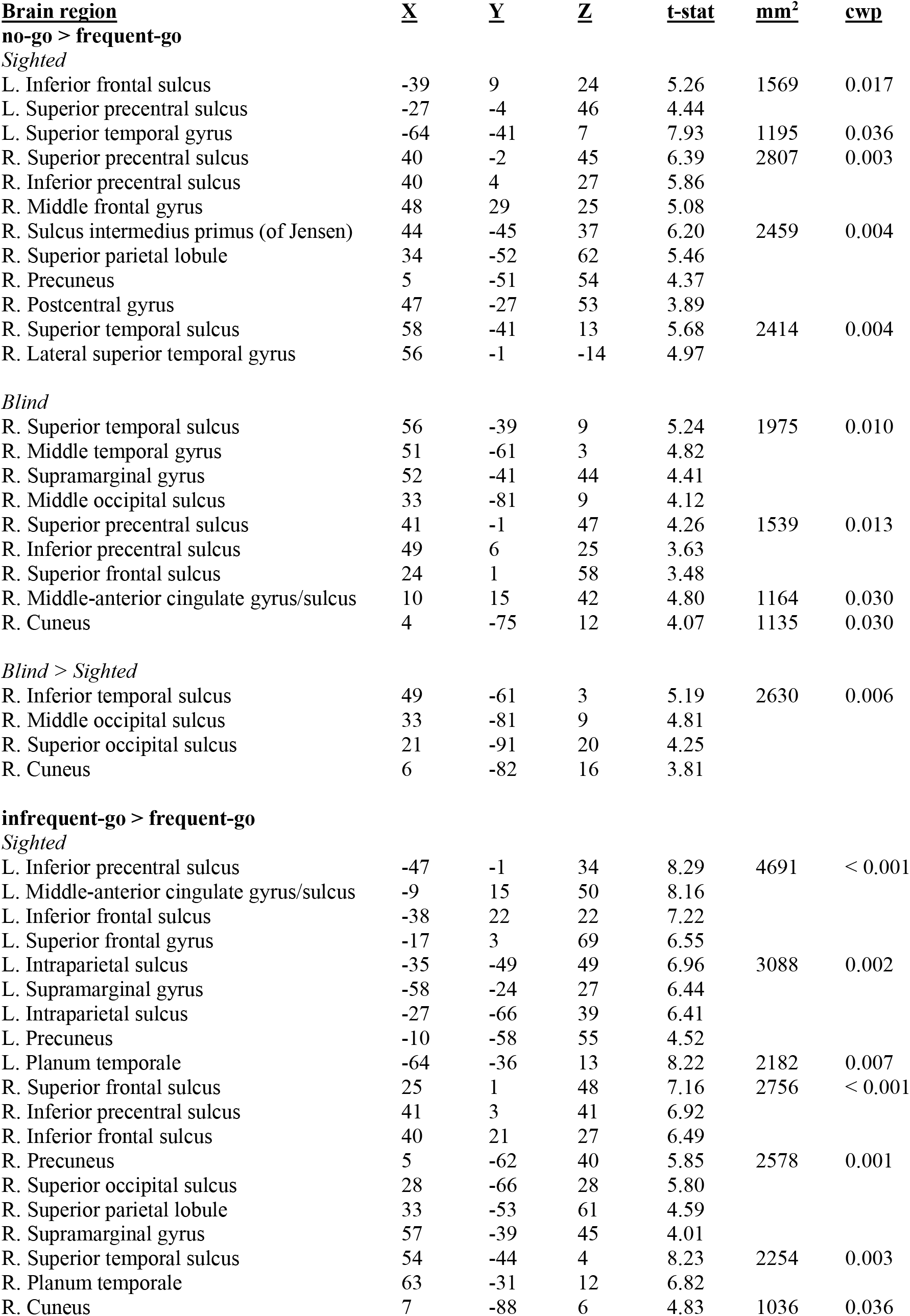
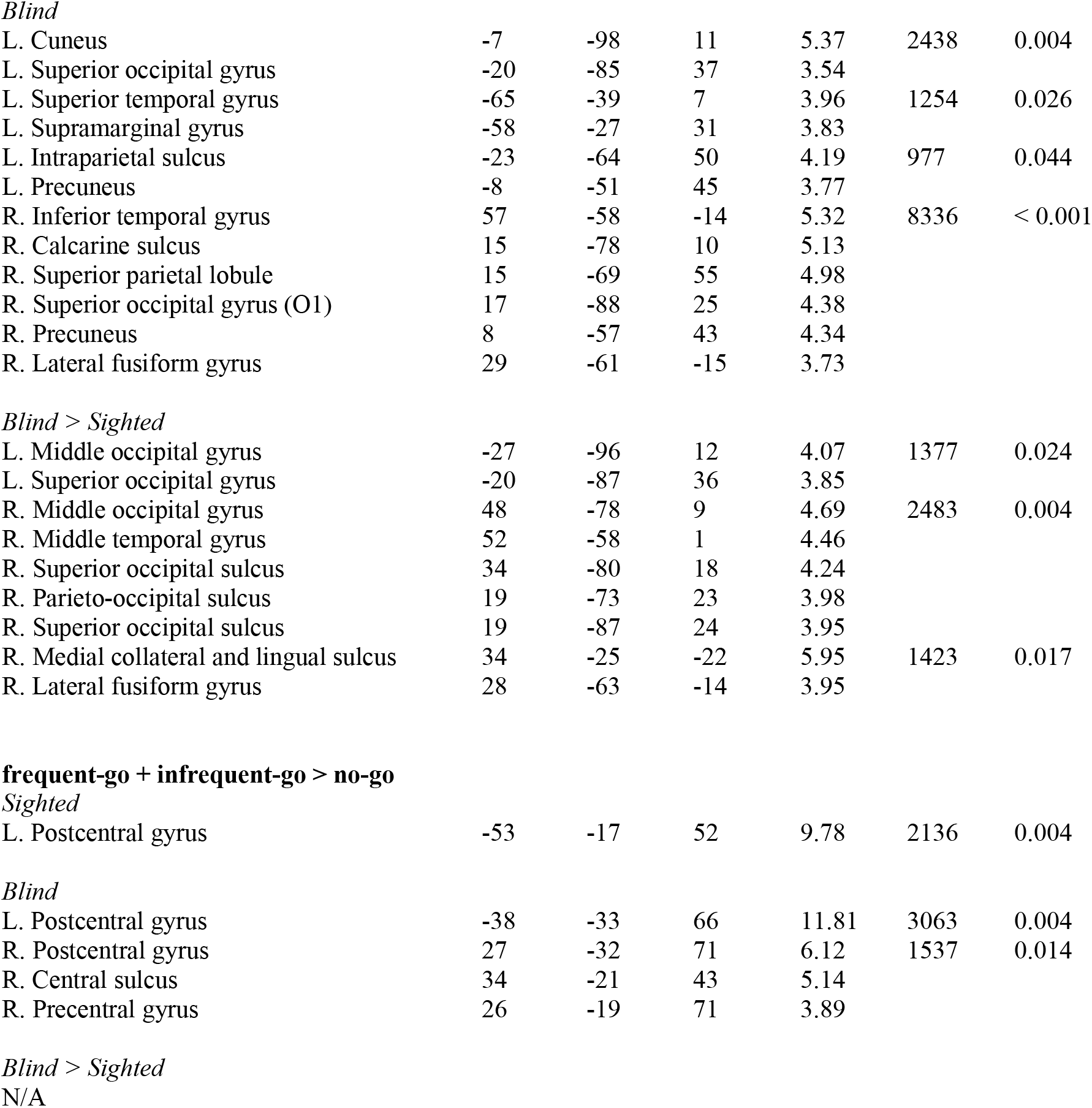
Brain regions differentially active across conditions, from cluster-corrected whole-brain analysis. Rows represent extrema, each characterized by a Destrieux Atlas gyral/sulcal name, an X- Y- and Z-MNI coordinate, and a t-stat(istic). Extrema are part of clusters, each of a mm^2^ size and a cluster-wise permutation probability. Extrema without listed cluster information are part of the preceding characterized cluster.

## Supplemental Results

We also looked for a sensorimotor response in the bilateral medial visual cortex (in case effects were lateralized, as in the sensorimotor cortices). Results were similar to those obtained in the right visual cortex. No sensorimotor effect was found in the bilateral “visual” cortices of blind participants (Supplementary Figure 1; frequent-go vs no-go t(18)=0.36, p>0.5; infrequent-go vs no-go t(18)=2.09, p=0.051; frequent-go vs. infrequent-go t(18)=2.26 p=0.03). In the sighted group, the visual cortices responded more to the two button press conditions (Supplementary Figure 1, frequent-go vs no-go t(18)=2.20, p=0.04; infrequent-go vs no-go t(18)=2.27, p=0.04; frequent-go vs. infrequent-go t(18)=0.06 p>0.5).

In bilateral V1 (across all vertices), we observed preferential activity for the infrequent-go condition within the sighted group (Supplementary Figure 1; infrequent-go vs frequent-go t(18)=2.45, p=0.03; infrequent-go vs no-go t(18)=1.73, p=0.10; no-go vs. frequent-go t(18)=0.08 p>0.5). In contrast, bilateral V1 of the blind group responded preferentially to both infrequent conditions (Supplementary Figure 1; infrequent-go vs frequent-go t(18)=3.42, p=0.003; no-go vs. frequent-go t(18)=2.89 p=0.01; infrequent-go vs no-go t(18)=0.26, p>0.5).

